# Measurement of Prostate Volume with MRI (A Guide for the Perplexed): Biproximate Method with Analysis of Precision and Accuracy

**DOI:** 10.1101/856542

**Authors:** Neil F. Wasserman, Eric Niendorf, Benjamin Spilseth

## Abstract

**Purpose:** To review the anatomic basis of prostate boundary selection on T2-weighted magnetic resonance imaging (MRI). To introduce an alternative 3D ellipsoid measuring technique that maximizes precision, report the intra- and inter-observer reliability, and to advocate it’s use for research involving multiple observers.

**Methods:** A demonstration of prostate boundary anatomy using gross pathology and MRI examples provides background for selection of key boundary marks when measuring prostate volume. An alternative ellipsoid volume method is illustrated. An IRB approved retrospective study of 140 patients with elevated serum prostate specific antigen levels and/or abnormal digital rectal examinations was done with T2-weighted MRI applying the new (Biproximate) technique. Measurements were made by 2 examiners, correlated with each other for interobserver precision and with an expert observer for accuracy. Correlation statistics, linear regression analysis, and tests of means were applied using p≤0.05 as the threshold for significance.

**Results:** Inter-observer correlation (precision) was 0.95 between observers. Correlation between these observers and the expert (accuracy) was 0.94 and 0.97 respectively. Intra-observer correlation for the expert was 0.98. Means for inter-rater reliability and accuracy were all the same (p=0.001).

**Conclusions:** Anatomic foundations for the boundaries of the prostate are reviewed. Precision and accuracy of total prostate volume using an alternative method is reported and found to be reproducible.

## Introduction

Measurement of the prostate gland *in vivo* is commonly required in the management of prostate disorders, both benign and malignant. Measurement is especially useful in the diagnosis and management of adenocarcinoma and benign prostatic hyperplasia (BPH). In the former, knowledge of total prostatic volume is necessary in the calculation of prostate specific antigen (PSA) density (PSAD)—a key indicator of the likelihood that elevated PSA is due to malignancy [1, 2]. Accurate measurement may affect cancer treatment strategies [1, 2]. However, the ability to make consistent and accurate measurement of the prostate is also useful in the diagnosis and management of BPH [3, 4].

Early attempts to estimate total prostatic volume by digital rectal examination (DRE) alone were replaced by more accurate measurement with transrectal ultrasound (TRUS) that showed excellent correlation with gross pathologic specimens using planimetric and calculated ellipsoid volume formula (EVF) techniques [5]. Similarly, accuracy compared to pathologic specimens has also been established using 3-plane magnetic resonance imaging (MRI) [6, 7, 8, 9]. While the forgoing techniques have shown reasonable success, details describing exact landmarks of the boundaries measured on imaging are rarely provided to assure general reproducibility. Difficulty in identifying landmarks such as the bladder neck and prostatic apex on ultrasound has been mentioned [10, 11]. Most studies performed with TRUS or MRI underestimate overall pathologic measurement of size by weight [6, 7, 12–14] although this may be due to inclusion of the seminal vesicles (SV), vas deferens (VD) stumps and periprostatic tissue in weighing the pathologic specimens [8]. Measured against an expert panel, *in vivo* measurement with TRUS overestimates post-operative small specimen size and underestimates large prostates [15, 16]. However other investigators using MRI EVF volumetrics report overestimation compared to specimen volume (corrected for density of 1.05 gm/cc) [9].

There are scant studies of inter-observer reliability for the measurement of total prostatic volume using standard methods [10, 11, 17]. TRUS intra-rater reliability using the ellipsoid formula was examined in one study and was 0.93 [12]. Bangma, et al. [18] compared TRUS step-section planigraphy with several geometric multiplane models and found the prolate ellipsoid to be comparable (r=0.89). Early application of MRI using the EVF showed acceptable precision and accuracy in one small study [6].

The purpose of this study is three-fold. We review detailed anatomy required to more confidently find measurement boundaries. We introduce a new multiplane EVF approach to make these measurements using MRI, and report results of inter-rater and intra-rater reliability with this technique.

## Materials and Methods

### Patient Selection

“This study was approved by our institutional Internal Review Board and Ethics Committee (protocol number 00003941) with a waiver of informed consent”. 140 patients were selected from a population of men undergoing MRI because of abnormal PSA, digital rectal examination or both. Ages ranged from 41-76 (mean, 64 years). They were selected by the senior radiologist (NFW) with over 40 years of experience who acted as administrator/expert. All MRI lobar classification types of BPH were represented in the selection [4].

### Study Design and Statistics

Two body imagers with 10 years of experience reading prostate MRIs were assigned the task of measuring total prostatic volume (EN, BS) using T2-weighted MRI on the 140 patient cohort. They were provided with a spreadsheet showing only the patient hospital identification numbers. After determining the total prostate volumes, these spreadsheets were returned to the administrator. Intra-rater reliability was tested only on the expert’s data. Re-measurement of prostate volume was done between 3 and 9 months after primary measures. Inter-rater reliability (precision) for continuous variables for volumes was analyzed for actual paired case-to-case correlation using the Pearson product moment and Lin’s Coefficient of Concordance [19]. Linear regression was calculated to find r^2^ values, Y intercepts, p values and 95% confidence intervals (CI). These were calculated from the open access statistical programs at the National Institute of Water and Atmospheric Research (NIAWA .com) and QI Macro Statistics^®^ (KnowWare International, Inc., Denver CO, USA). Measurements of the administrator were used as a proxy for gross pathological specimen weight for the determination of rater “accuracy”. Correlations were analyzed by linear regression and graphed. A comparison of means was also done after the data was available to establish data normality. Significance was defined as p <0.05.

### MRI Acquisition Techniques

Prostate MRI was performed at 3T with a pelvic phased array coil in 27 with and 113 without endorectal coil (Magnetom Skyra or TrioTim; Siemens Healthcare; Erlangen Germany). T2W FSE imaging was performed in the axial plane (TR/TE 3700/80 msec; NEX 3; 3mm slice thickness; no interslice gap; flip angle 160 deg; FOV 140 mm, matrix 320 x 256) and coronal plane (TR/TE 4030/100 msec; NEX 2; 3 mm slice thickness; no interslice gap; flip angle 122 deg; FOV 180 mm; matrix 320 X 256). T2W 3D SPACE images were obtained (TR/TE 1400/101 msec; flip angle 135 deg; 256 x 256 x 205 matrix, 180 mm FOV). In addition to 1 mm axial images, T2W 3D SPACE images were reconstructed at 3 mm in the axial, sagittal, and coronal planes. Additionally, dynamic contrast enhanced T1-weighted (3 mm slice thickness; TR/TE 4.9/1.8 msec; 224 X 156 matrix, 250 mm FOV, temporal resolution <10 sec), and diffusion-weighted images at b values of 50, 800, and 2000 s/mm3 were performed.

### Anatomy

Anatomic boundaries and landmarks for measurement of total prostate volume were chosen after a thorough review of the anatomic literature, including the work of Salvadore Gil-Vernet (salvadoregilvernet.com) [20], Robert Meyers [21, 22] and others [23, 24, 25]. The Prostate is not only a glandular but also a muscular organ [23, 26]. Its external boundary, that may be referred to as the external prostatic capsule (EPC) extends posterior, lateral, and antero-lateral. The EPC represents the peripheral compressed stromal (fibro-muscular) component of the latticework supporting the glandular elements [23]. This is the outer boundary that is visible on imaging. Walz, et al. [27] refers to this as the “true capsule”, although it is not what some anatomists would define as a true “capsule” that can be “peeled off” and separated from the gland. It will be referred to as a “capsule” in this presentation due to its common use in clinical practice and because it is the outer boundary we see on imaging.

The controversy surrounding the naming of this “pseudocapsule” is discussed in several anatomic studies including those that refer to the periprostatic fascia as the “capsule” [27 25]. This fascial plane is not visible on imaging. The anterior external limit of the prostate is less distinct, comprising anterolateral elements of the EPC, the surgical capsule (SC) surrounding enlarged transition zones (TZ) when BPH is present, and anterior fibro-muscular stroma (AFMS). The multilayered loose fascia surrounding the prostate is disrupted with surgical or post-mortem removal making correlation between the *in vivo* gland to the *ex vivo* specimen more difficult [22].

The “surgical” capsule, (SC) is so-called because it encases the enlarged transition zone, which on open surgery can enucleated from surrounding tissues. It represents compressed circular smooth muscle fibers of the pre-prostatic urethral wall (above the verumontanum) and prostatic urethral wall (below the veru) [23]. The anterior SC is indistinct to absent unless there is more advanced nodular hyperplasia of the (TZ). The AFMS is thickest proximally where it is formed by smooth muscle fibers from the bladder wall. This boundary is critical to define when performing maximal AP prostate measurement. The above features are delineated in figures (1A-C). The EPC is incomplete at the apical peripheral zone (PZ) [22, 23, 24] (Fig. 1D).

**Fig. 1A.**
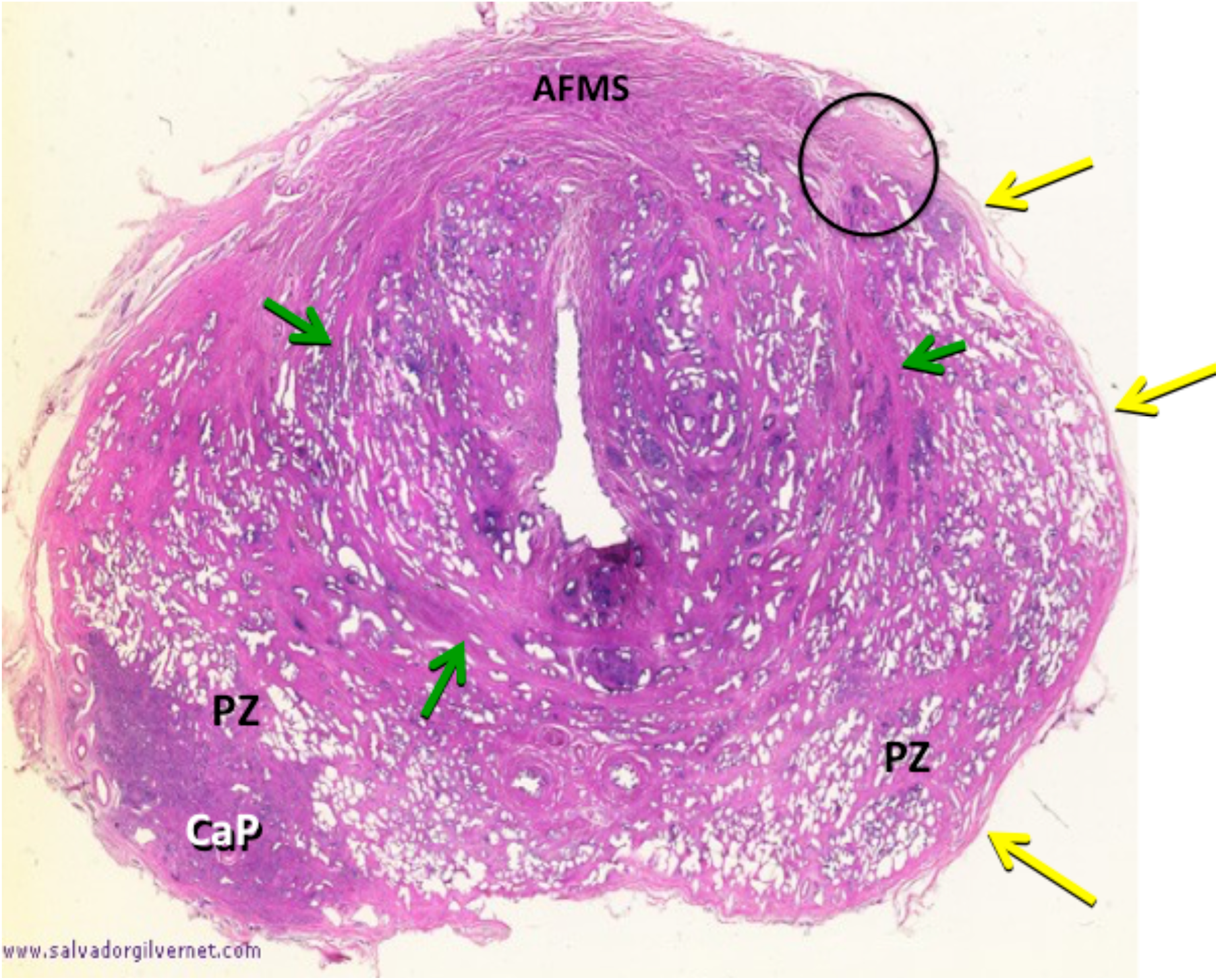
H & E stained transverse section of prostate with minimal TZ hyperplasia demonstrating the minimally expanded compacted outer circular smooth muscle fibers forming a surgical capsule (green arrows) and peripheral external prostatic capsule (yellow arrows). Notice how they fuse into the anterior fibromuscular stroma (AFMS.) PZ = peripheral zone, CaP = carcinoma of the prostate.) (Modified courtesy of José Maria Gil-Vernet, M.D.)

**Fig. 1B.**
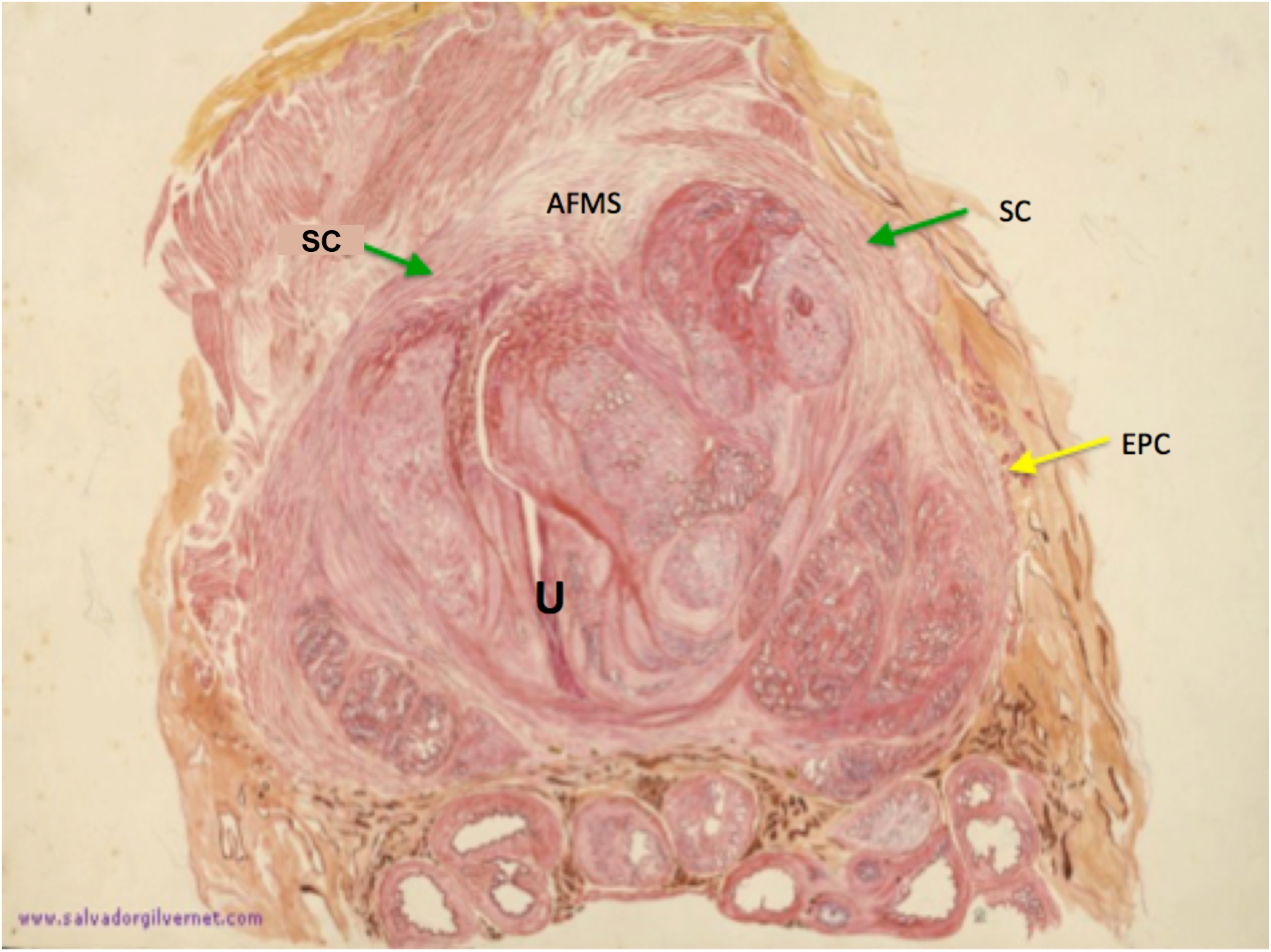
Transverse H & E stained section in a patient with advanced nodular hyperplasia of the TZ demonstrates the anterior “fused” EPC and SC on the left (yellow and green arrows respectively) which are incomplete anteriorly as is their counterpart on the right. The space in between is filled in by the anterior fibromuscular stroma. External prostatic capsule (yellow arrows), Surgical capsule (green arrows). U = urethra, AFMS = anterior fibromuscular stroma. (Modified courtesy of José Maria Gil-Vernet, M. D.)

**Fig. 1C.**
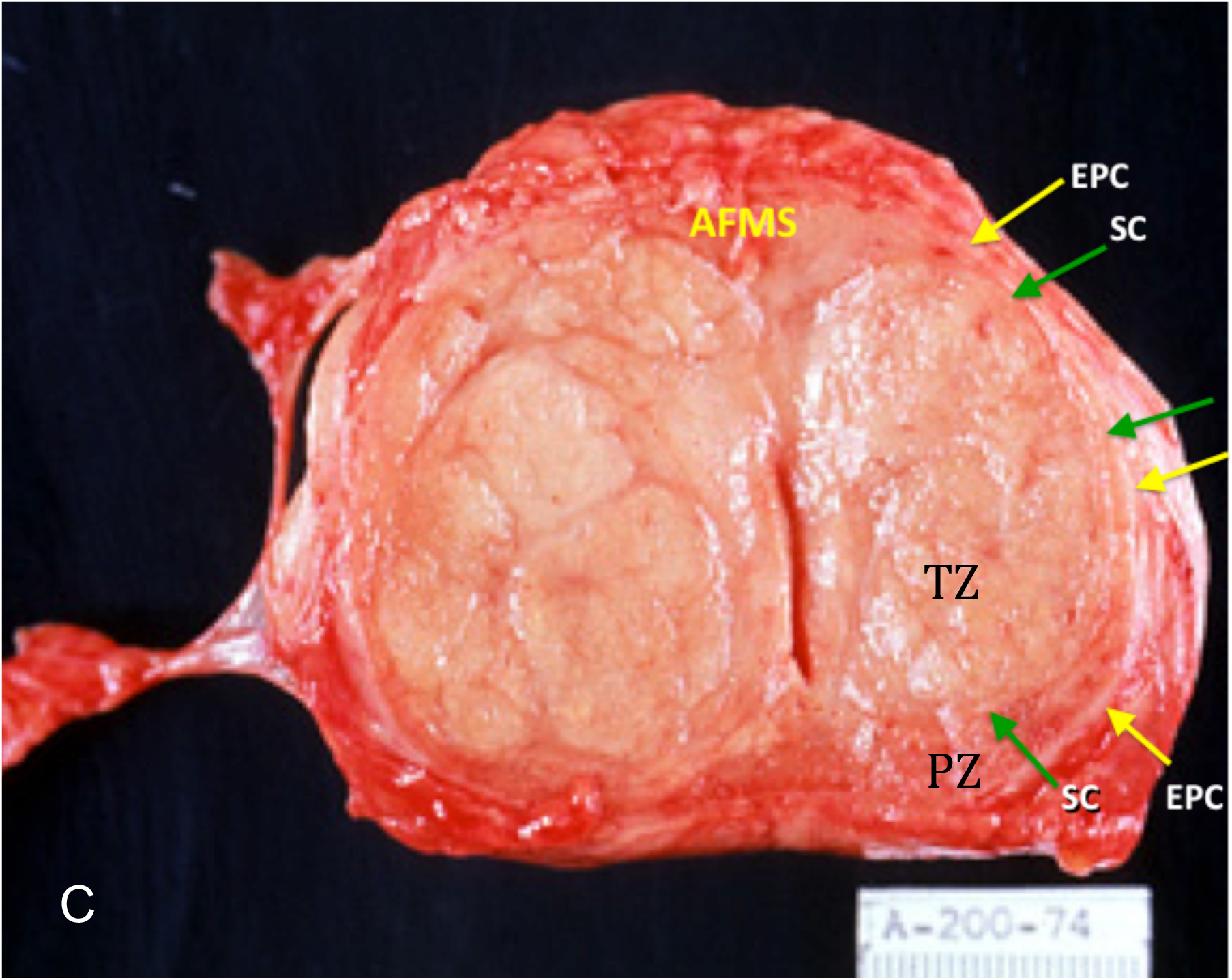
Gross specimen of patient with advanced BPH showing EPC (yellow arrows) and SC (green arrows) merging anteriorly with the anterior fibromuscular stroma (AFMS).

**Fig. 1D.**
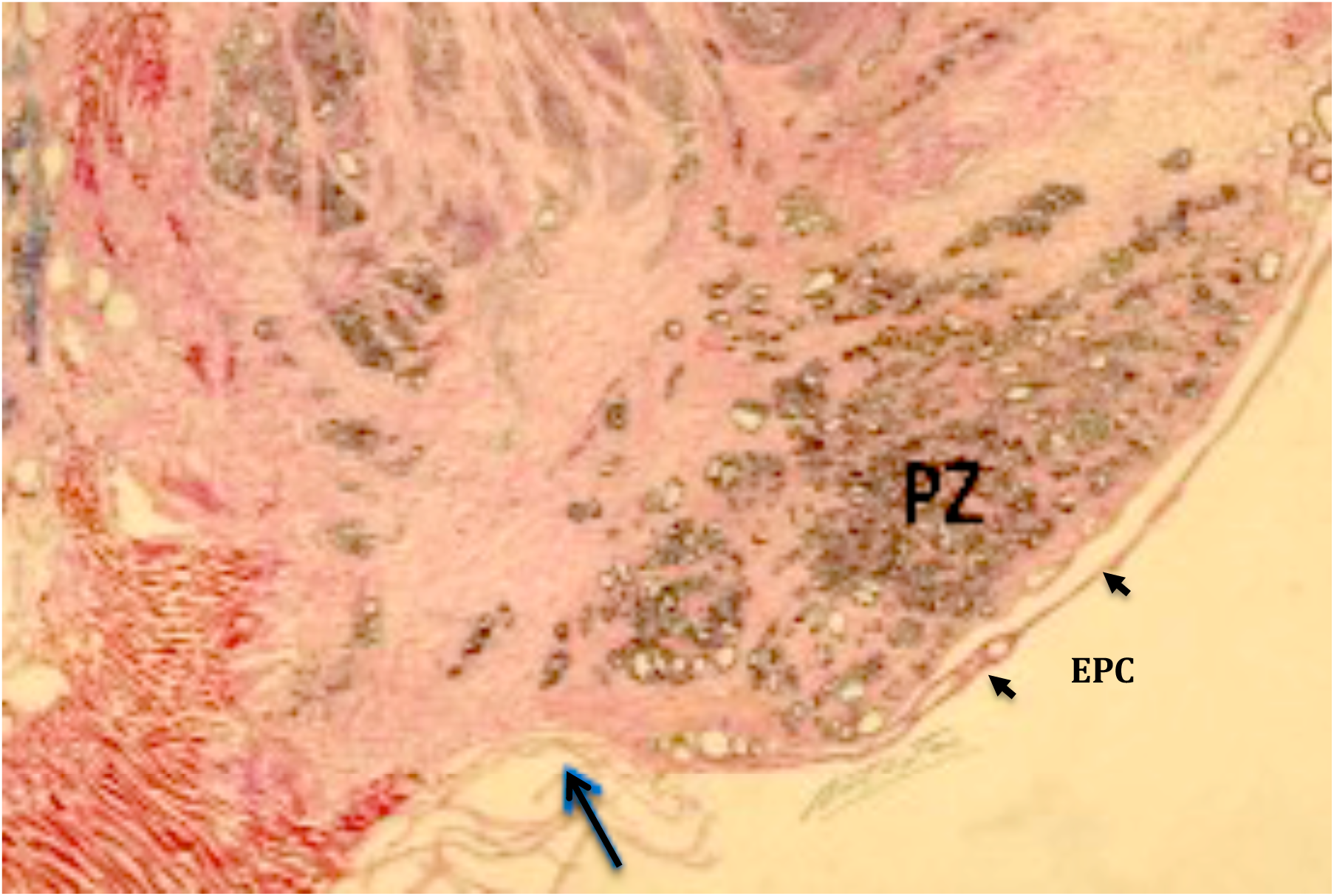
Parasagittal low magnification H & E section showing absence of the EPC at the apex (arrow)

### MRI Image Analysis & Selection of Boundaries

All images were reviewed on high-resolution monitors using Philips Intellispace (Koninklijke Philips N.V.) software. Because of new interest in the urological literature pertaining to herniated portions of enlarged prostate into the bladder base, an alternative scheme of volumetrics was devised to include measurement of intravesical prostatic protrusion (IPP). A new vocabulary is needed to describe these boundaries e.g. vesico-prostatic angle (VPA), vesico-prostatic line (VPL), and apical line (AL) discussed below. Our approach is calibrated to provide the most reproducibility for intra-and interrater reliability.

Transverse measurements were made in the axial plane estimated to show maximal diameter. It allows best visualization of the lateral EPC boundaries for transverse measurement. The axial plane is also the only plane that allows for selection of the anterior boundary by interpolation of anterior ends of the right and left EPC for accurate antero-posterior (AP) measurement. The mid-sagittal plane is unsatisfactory for that determination. (The coronal plane, though reasonable for transverse measurement, is suboptimal for AP diameter measurement.)

Pericapsular veins and levator ani muscles are confounding anatomical structures that, depending on axial level, lie adjacent to the EPC and make its outer boundary difficult to define (Fig. 2A). Therefore, for consistency, when measuring the transverse diameter, we arbitrarily chose to measure from the inner margins of the EPC.

**Fig. 2A.**
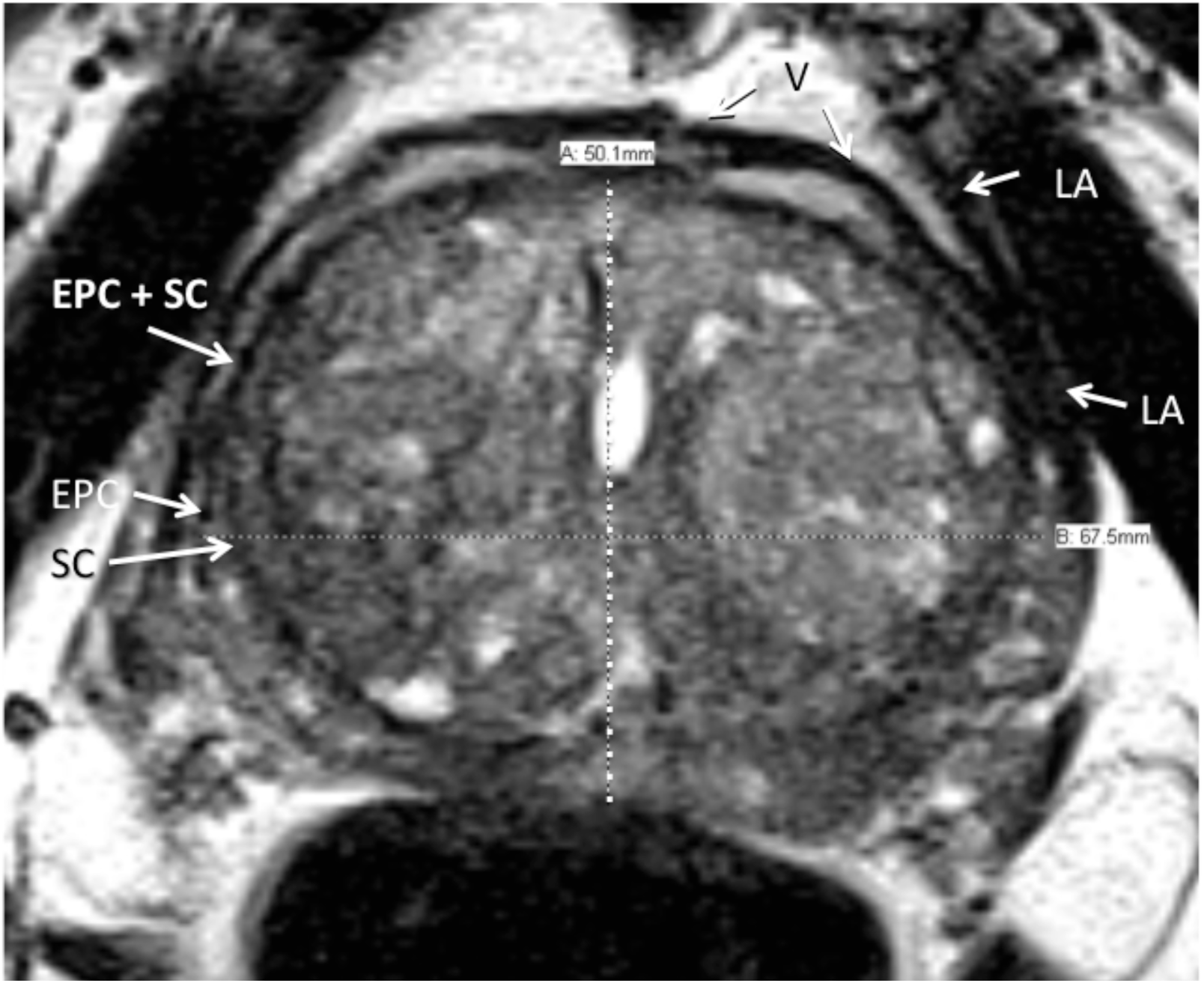
Axial T2-weighted axial MRI. Structures surrounding the prostate that might be mistaken for the external prostatic capsule (EPC). These include the periprostatic veins (V) and levator ani muscle (LA).

**Fig. 2B.**
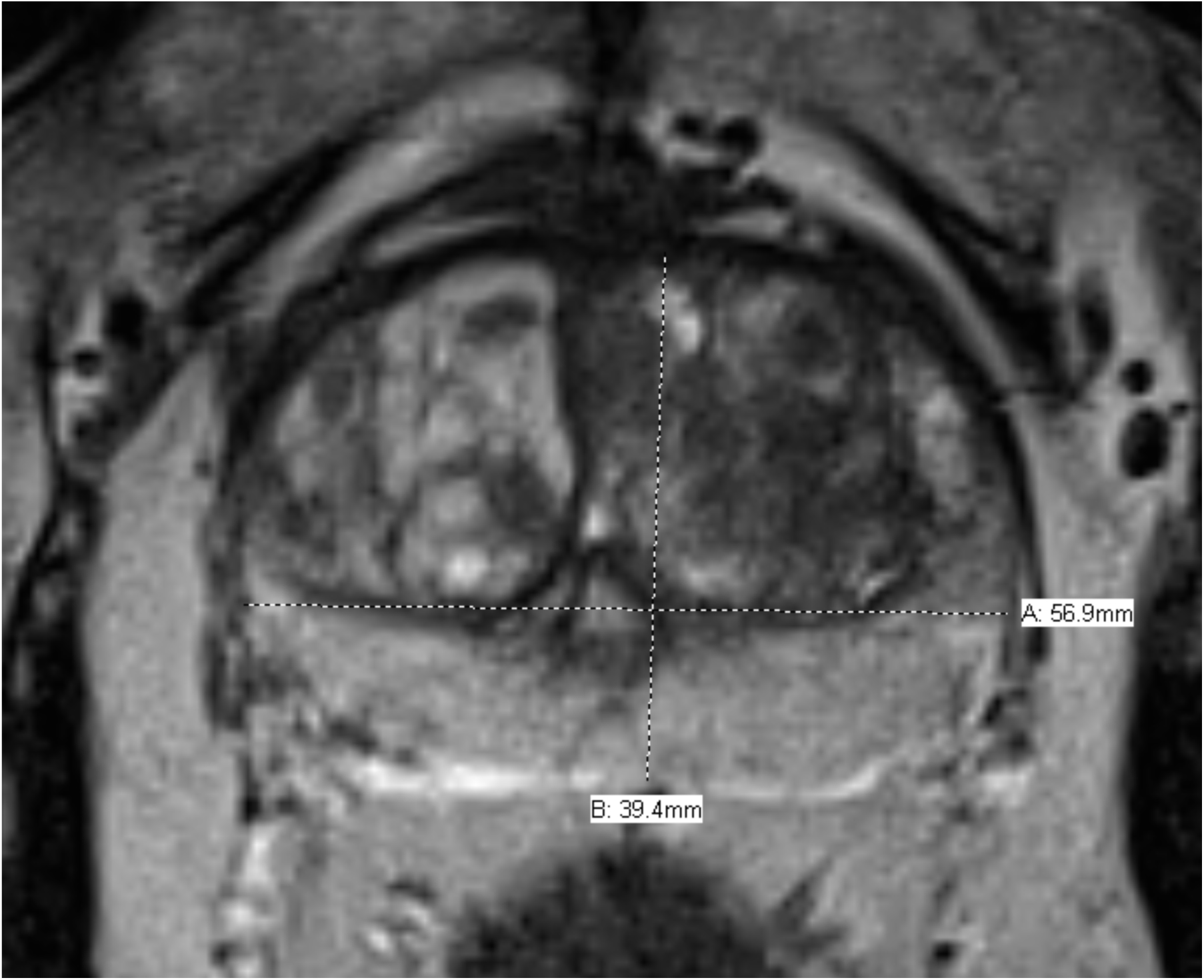
Axial T2-weighted axial MRI. Transverse measurement is arbitrarily made from inside border of external prostatic capsule (EPC).

The maximal AP diameters were made at or near the same axial plane as the transverse taking care to not include the inferior extension of the anterior bladder detrusor. Measurement was made from the inner margin of the mid-posterior EPC to the mid-anterior AFMS. If there was a midline posterior indentation of the EPC (posterior commissure), the inner point was selected, even though the PZ projected slightly more posterior on either side. The anterior landmark chosen varied depending on definable anatomy.

There are three general configurations of the anterior boundary. In some patients, there is a discrete continuation of the merged SC and bilateral anterolateral EPC across the midline allowing for selection of its outer boundary (Fig. 3A). In the second, the EPC is interrupted and invisible in the midline, in which case we drew an interpolated arc-like line connecting the ends of these structures (Fig. 3B). (This is the rationale for using the axial rather that sagittal plane for AP landmarks.) The third configuration is where there is apparent complete fusion of the AFMS, SC and EPC, completely obliterating visualization of the anterior EPC. This configuration was seen in less than 5% of our cohort. An interpolated arc-like line was dawn if the two antero-lateral ends of the EPC were close enough to connect. Otherwise, the axial AP anterior landmark was taken as the inner boundary of the AFMS (Fig. 3C). The appearance may be caused by the “salami effect” of a trans-axial plane that is oblique with respect to the longitudinal axis of the prostate (Fig. 4).

**Fig. 3A.**
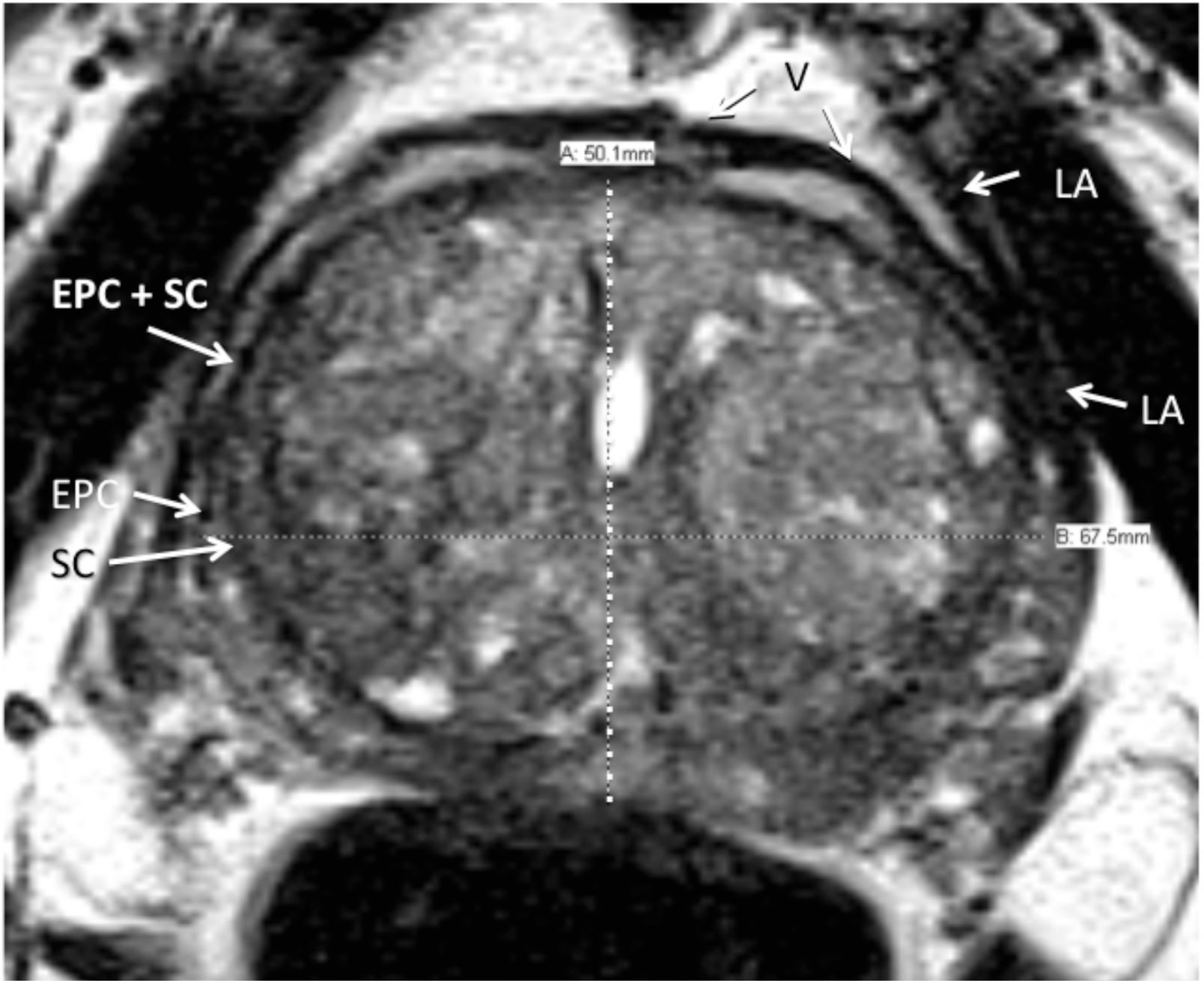
Axial T2-weighted axial MRI. Anterior boundary is clearly present at mid-sagittal conjunction of right and left EPC.

**Fig. 3B.**
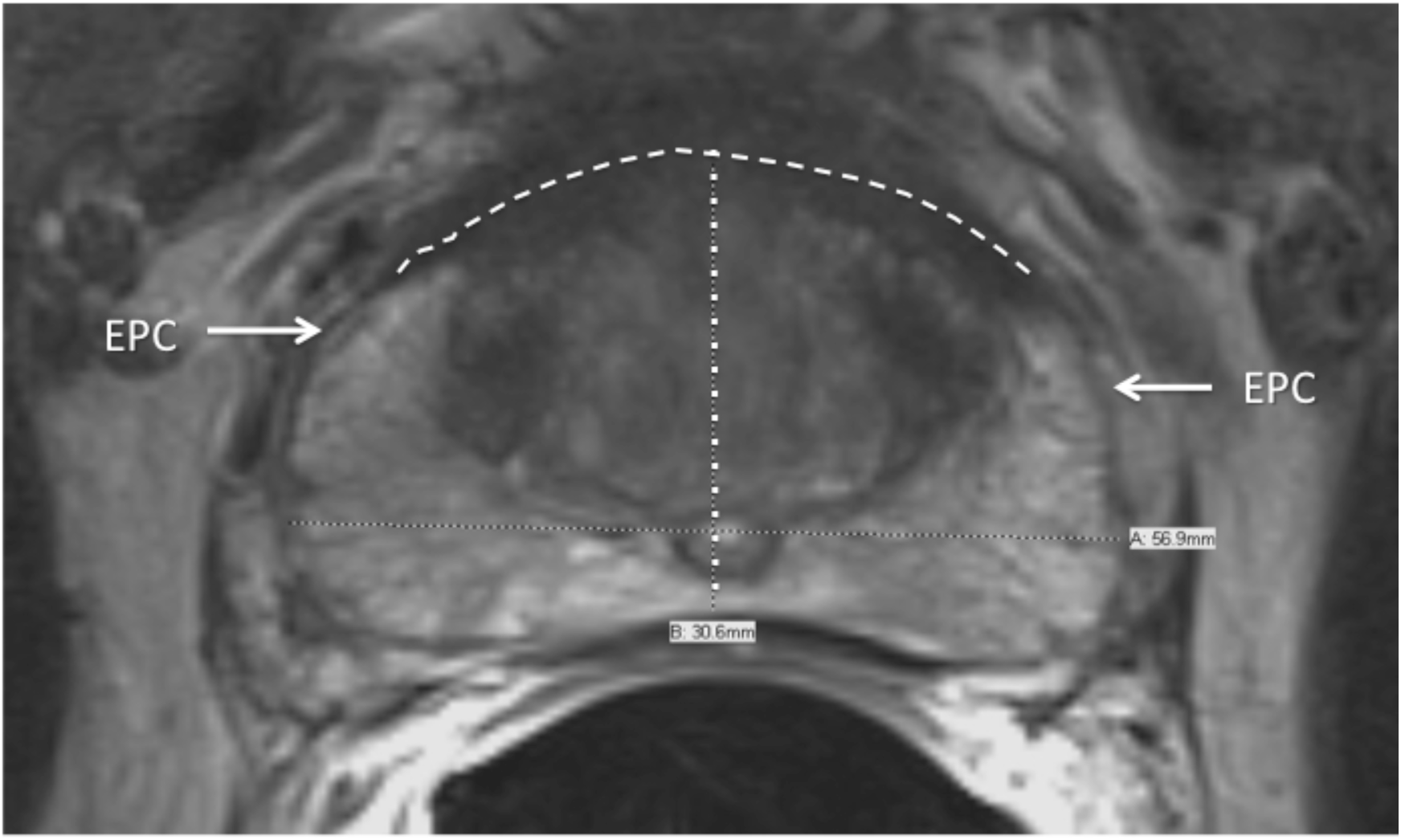
Axial T2-weighted MRI. Interpolated Arched line connecting anterior free ends of right and left EPC. Dotted line is antero-posterior mesurement

**Fig. 3C.**
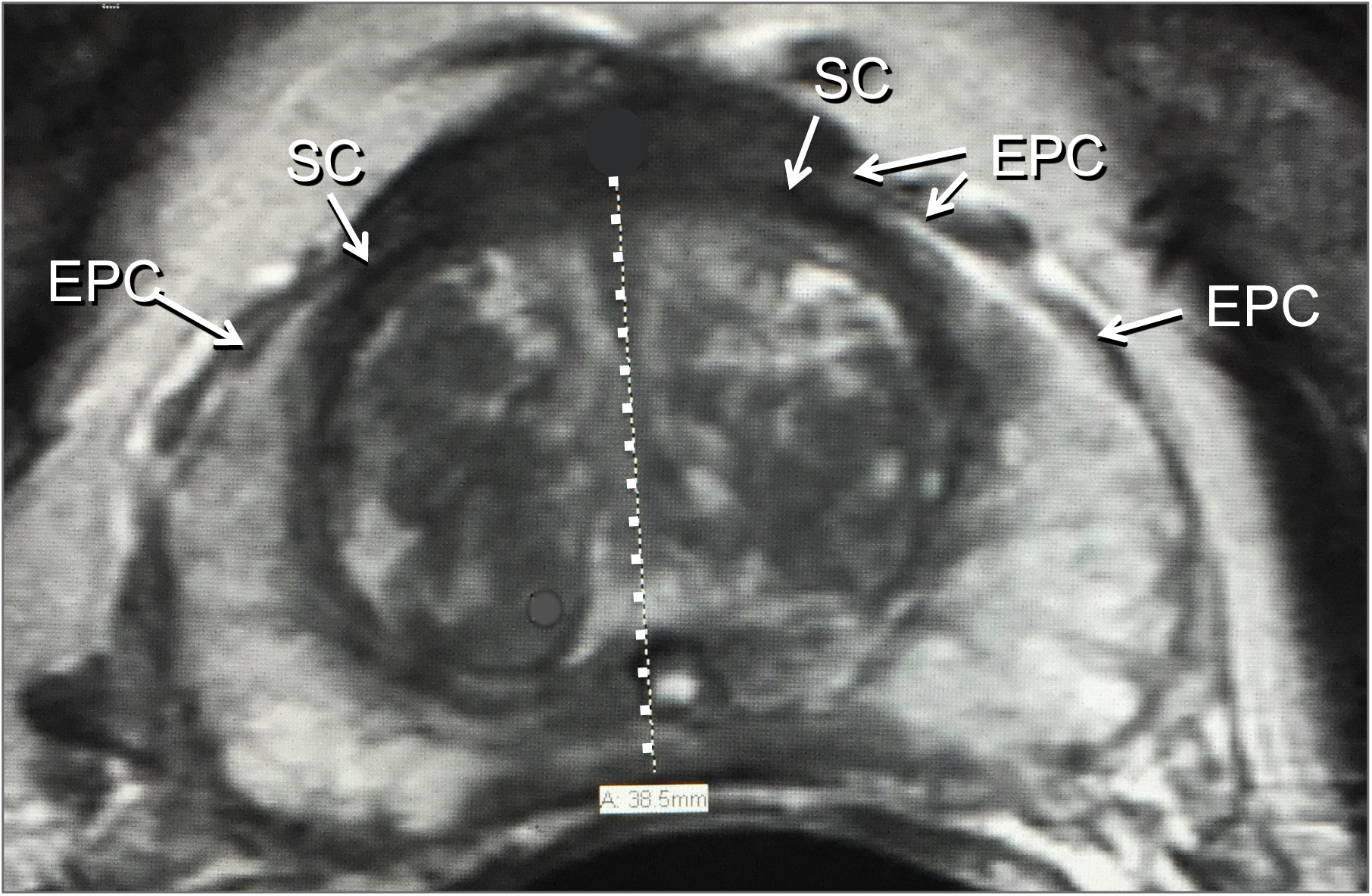
Axial T2-weighted MRI. Right and left external prostatic capsule (EPC) merge into anterior fibromuscular stroma. SC=surgical capsule

**Fig. 4.**
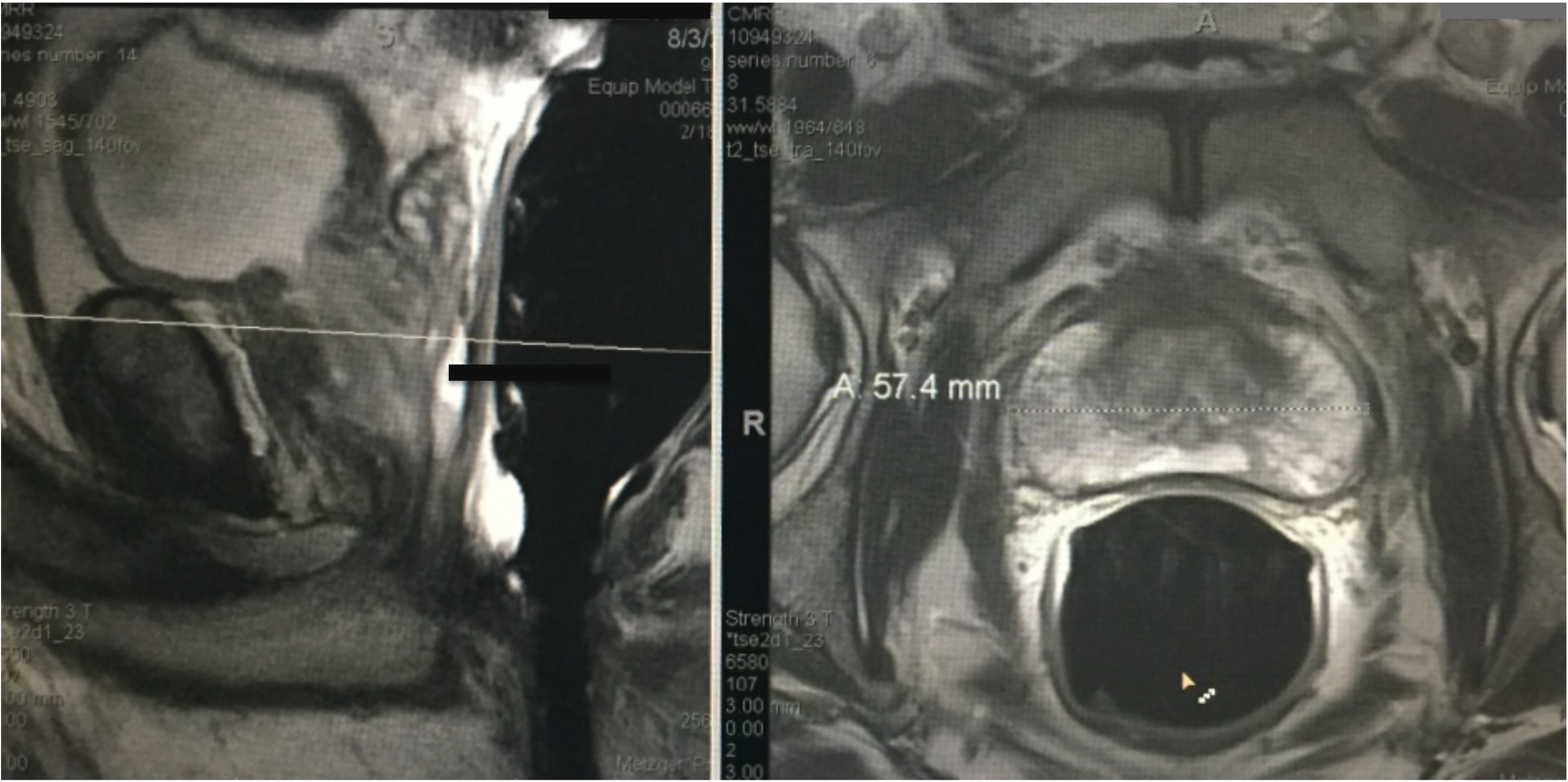
T2-weighted MRI in Mid-sagittal plane (left image) with transaxial line oblique with respect to longitudinal axial line from bladder outlet to apex produces “Salami effect”. This results in a non-orthogonal anterior plane through the proximal thickest portion of the anterior fibromuscular plane (right image). This case is not from our cohort since it shows use of endorectal coil.

Length measurement was made in the mid-sagittal plane. The mid-sagittal plane is defined as that which best shows the membranous and bulbous urethra. Landmarks for measuring mid-sagittal length started by finding the anterior and posterior vesico-prostatic angles (VPA) as defined as the points where the detrusor connects with the prostate and connecting these angles by a line called the vesico-prostatic line (VPL) (Fig. 5). The VPL arbitrarily defines the level of bladder “outlet.” The inferior landmark was the caudal margin of the PZ defining the apex where an apical line (AL) was drawn approximately perpendicular to the long axis of the prostate drawn from the bladder outlet to the membranous urethra (Fig. 5). This PZ inferior boundary may occur posteriorly or in a parasagittal plane, therefore, it is essential to use a localizer tool in the coronal view to verify the level of the AL in the mid-sagittal plane (Fig. 6). If the prostate was entirely below the VPL, the mid-sagittal length was defined as the length of a line drawn from the mid-urethra at the level of the AL extending perpendicular to the VPL. If the prostate projected above the VPL, the midsagittal length was defined as the sum of the distance below the VPL plus the distance of a line from the from the superior margin of this tissue drawn perpendicular to the VPL (Fig. 5). This segment superior to the VPL represents intravesical prostatic protrusion (IPP).

**Fig. 5.**
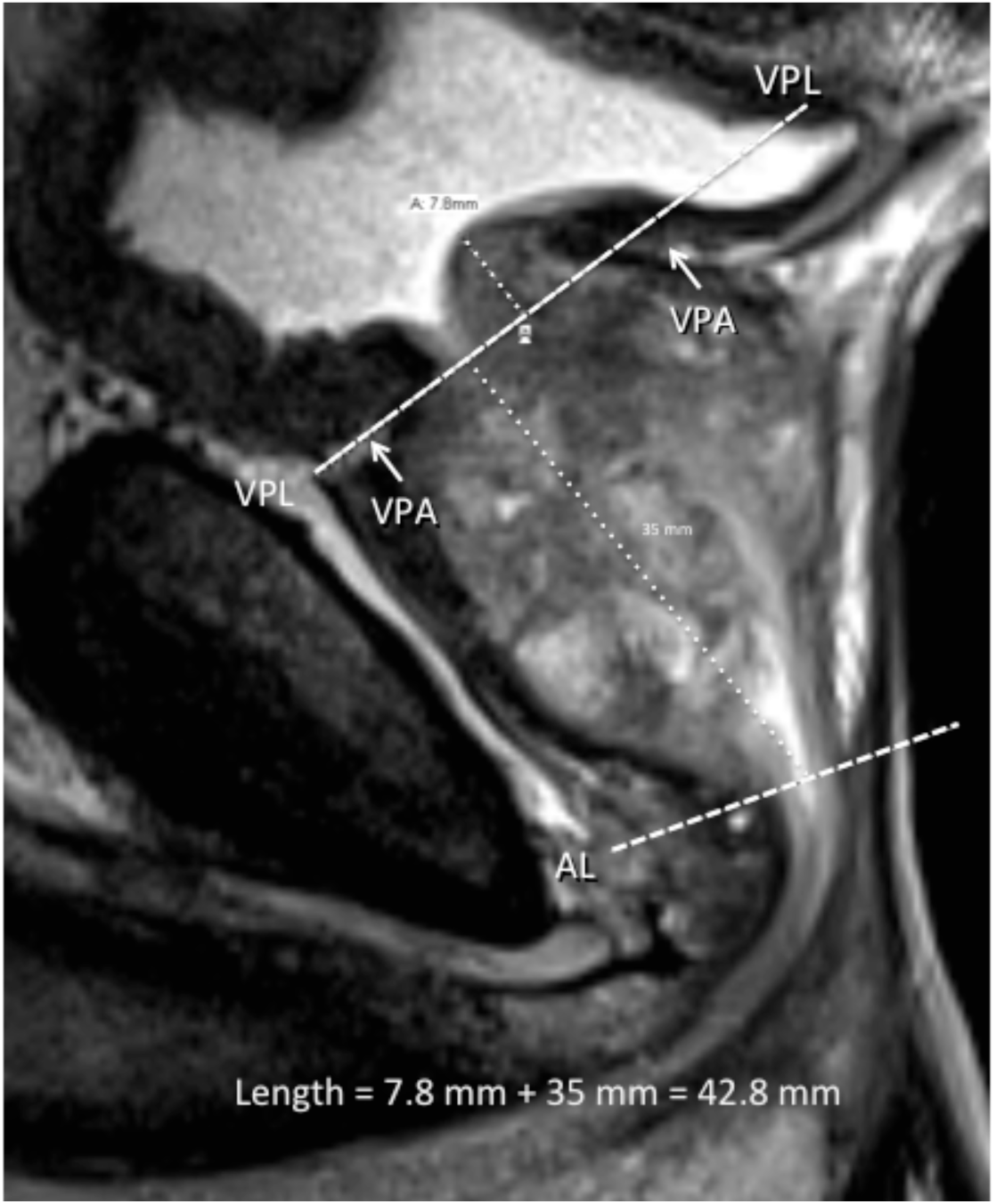
Midsagittal T2-weighted MRI showing the vesico-prostatic angles (VPA) and drawing the dashed vesico-prostatic line (VPL). The dashed apical line (AL) is also demonstrated. Dotted lines represent length measurements summed to establish length of prostate for use in prolate ellipsoid formula to determine total prostate volume.

**Fig. 6.**
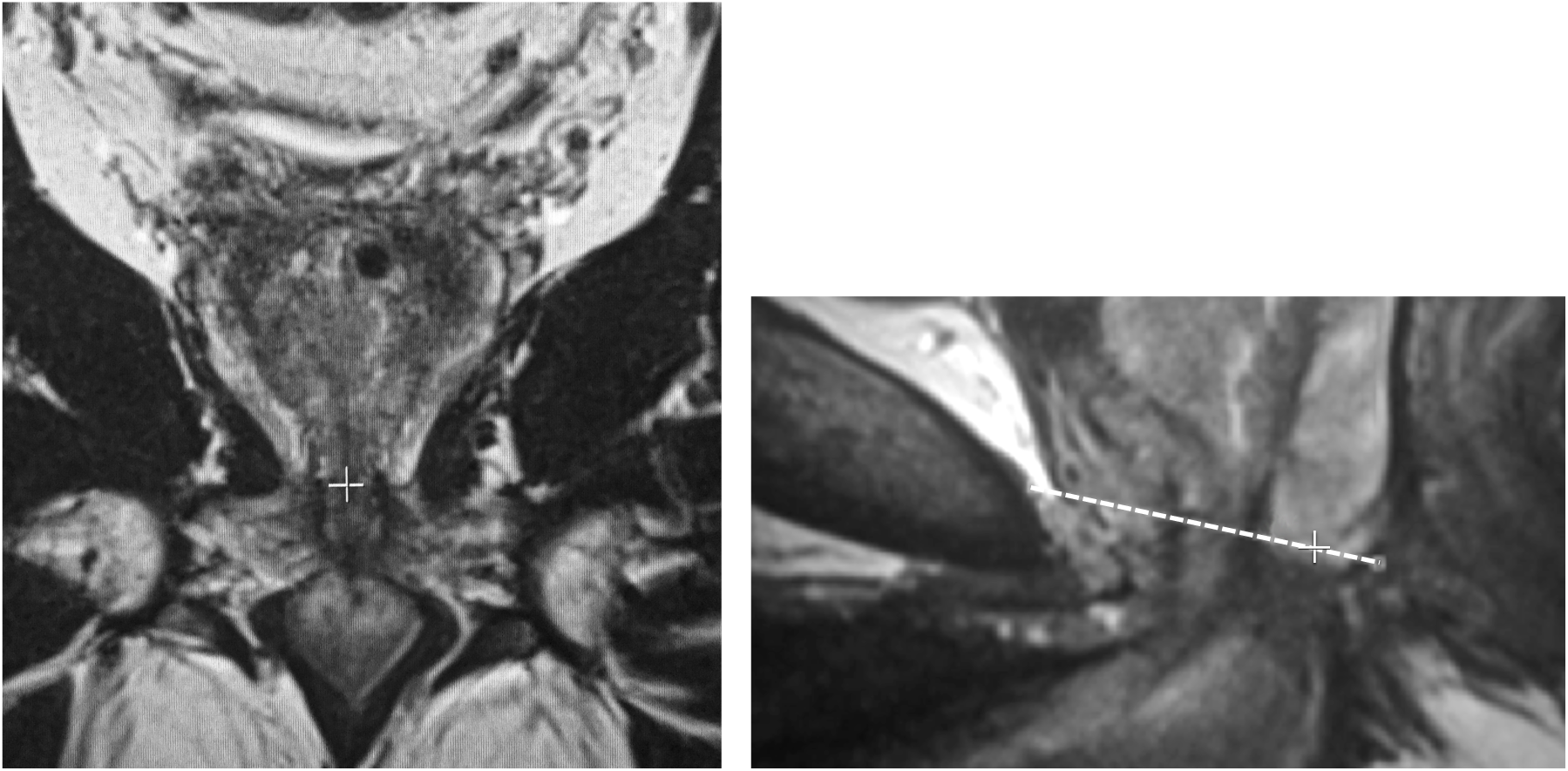
T2-weighted MRIs. Accurate apical localization requires use of the coronal plane. The apex, as defined as the inferior boundary of the peripherl zone (PZ) is located in the coronal plane (+ on left image) to simultaneously establish the correct level of the apex in the mid-sagittal plane (right image) where + sign is superimposed over the membranous urethra

**Fig. 6B.**
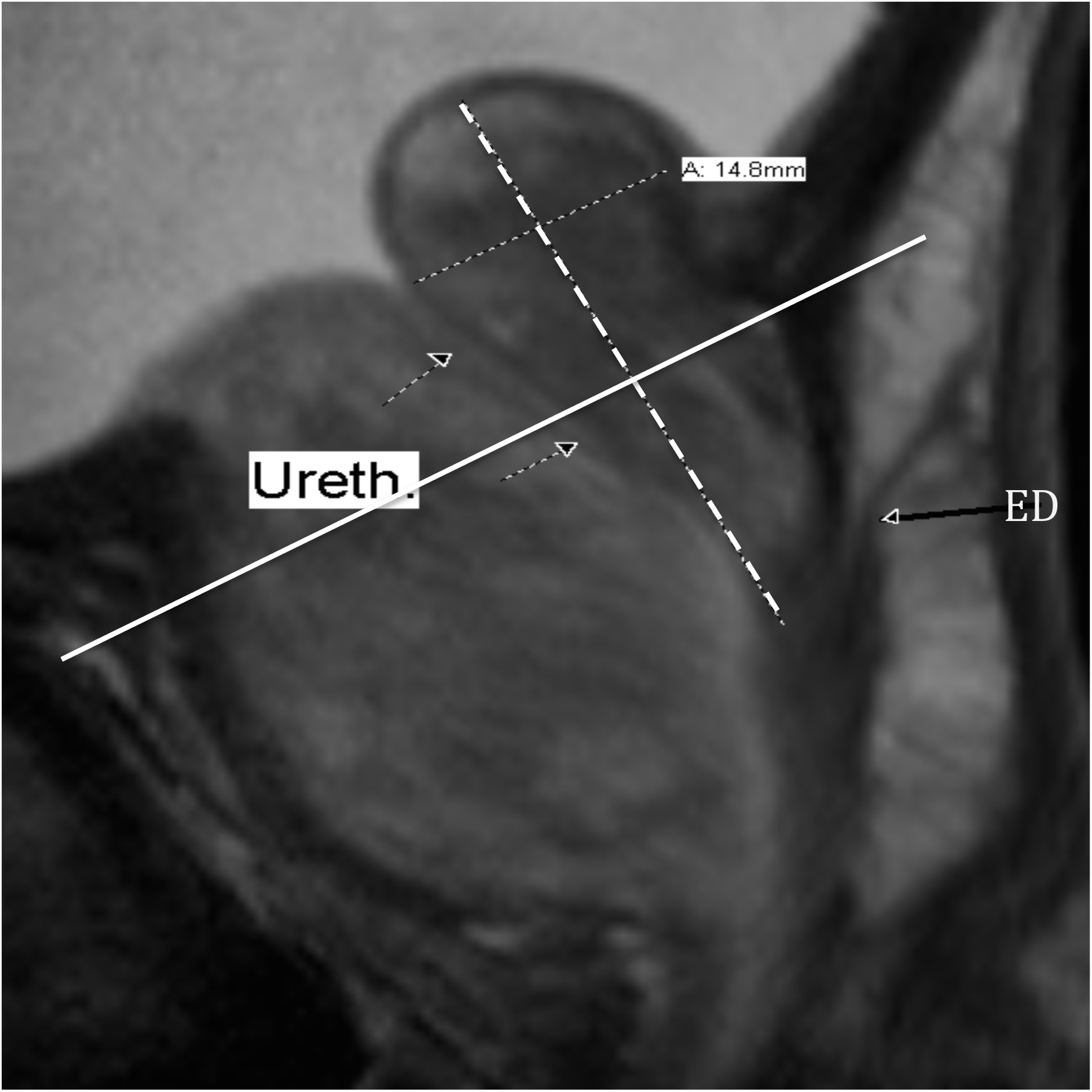
Mid-sagittal T2-weighted MRI. Type 3 benign prostatic hyperplasia where maximal antero-posterior (AP) diameter is above the vesico-prostatic line (VPL). Dashed line shows the long axis of the retro-urethral lobe. The large dotted freehand line marks the course of the urethra. In this case, the AP measurement must be above the VPL (solid line). Ejaculatory duct entering the upper verumontanum (ED) is indicated.

Total prostatic volume was calculated according to the formula of the geometric model for a prolate ellipsoid: Volume = AP x Transverse x Length x π/6

The size of the retrourethral lobe (RU) was arbitrarily evaluated by measuring the maximal AP diameter as visualized on the mid-sagittal view. This involved three steps: Step 1: The RU long axis was defined by a line from the cranial margin of the RU to the mid-verumontanum (conjunction of the ejaculatory ducts with urethra). Step 2: We traced the approximate course of the urethra from the bladder neck through the level of the verumontanum using the “freehand” tool. Step 3: we estimated the thickest AP level of the RU as measured from the inner margin of its posterior capsule perpendicular to its long-axis ending at the posterior wall of the urethra (Fig. 7A, B).

**Fig. 7A.**
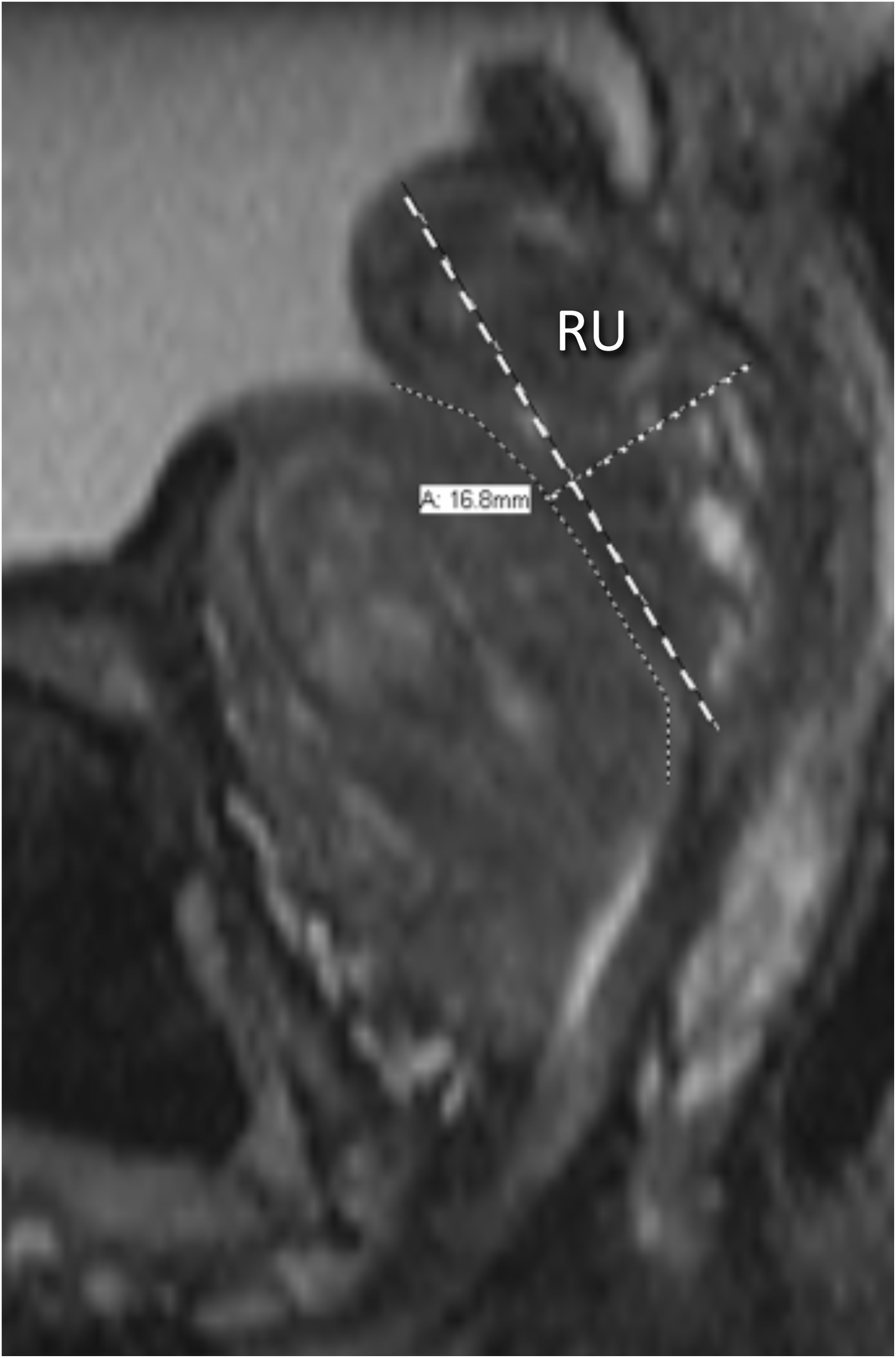
T-2-weighted mid-sagittal MRI showing the 3-step measurement of the r etrourethral lobe (RU). The anterior enlarged transition zone (TZ) displaces the urethra posteriorly (freehand dotted line). Type 3 BPH with a high intra-vesicle retrourethral lobe (RU) is protruding into the bladder (IPP). The long axis of the retrourethral lobe (dashed line) extends from cranial boundary of the RU to the verumontanum. The final step in the antero-posterior measurement of the RU is a line perpendicular to the long axis from the posterior RU boundary to the urethra (large dotted line). In this case, we suggest using a mid-gland AP measurement, because it was greater than the one above the VPL.

## Results

Analysis of the alternative method of determining total prostatic volume resulted in an interobserver correlation (precision) of 0.95 for the two interpreting radiologists (Fig. 8A). Intra-observer correlation (precision) was 0.98 for the senior radiologist (administrator) (Fig. 8B). Interobserver correlation between the two interpreting radiologists and the administrator (accuracy) was 0.94 and 0.97 respectively (Fig. 9A, B). All correlation p values <0.000. Since the data was non-normal in distribution, a parametric test of total prostate volume means (Wilcoxan Sign Rank) was applied and means for interrater reliability (precision) and accuracy compared to the administrator were all the same p=0.001 (Fig. 10].

**Fig. 8A.**
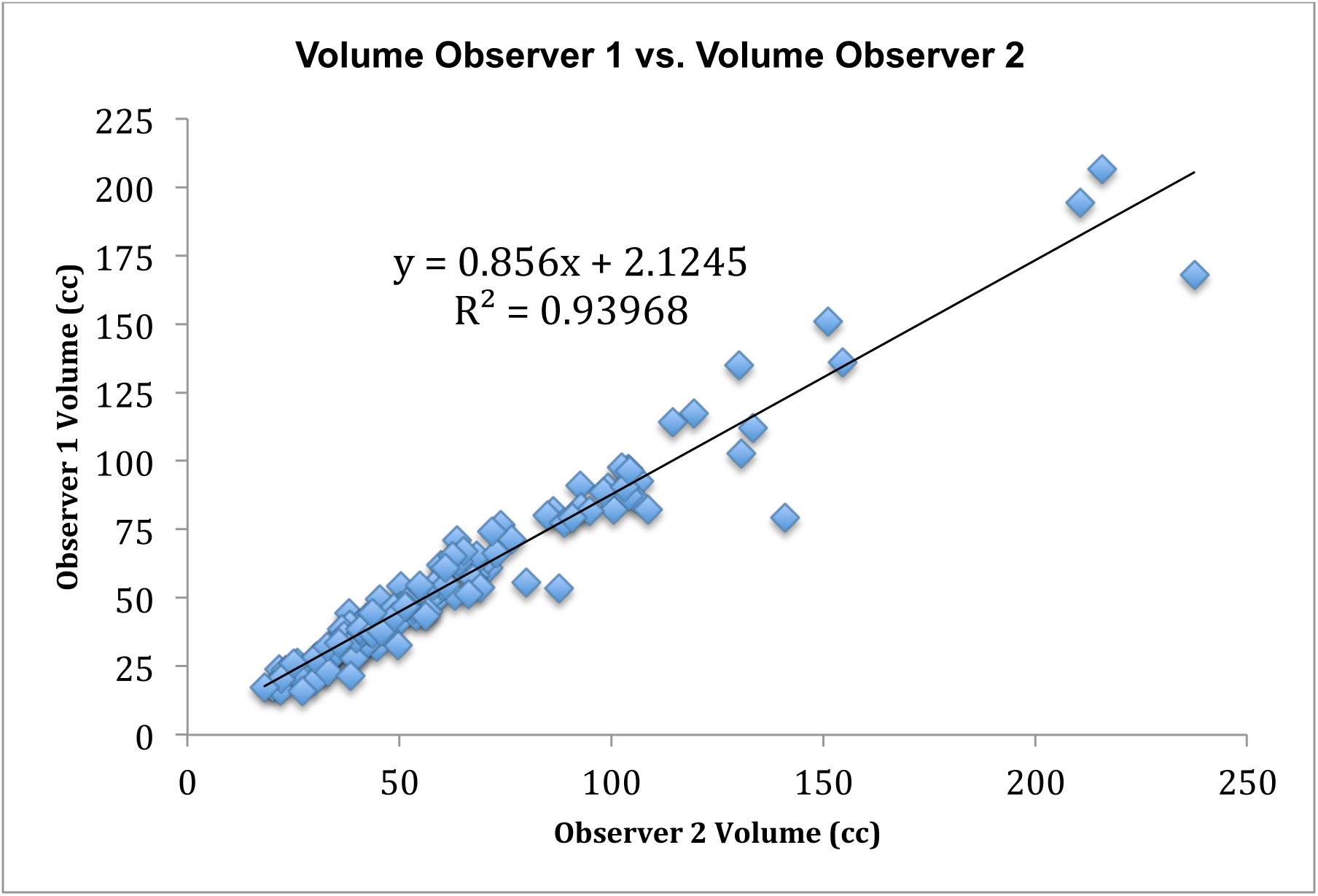
Inter-rater Concordance Between Two Interpreters

**Fig. 8B.**
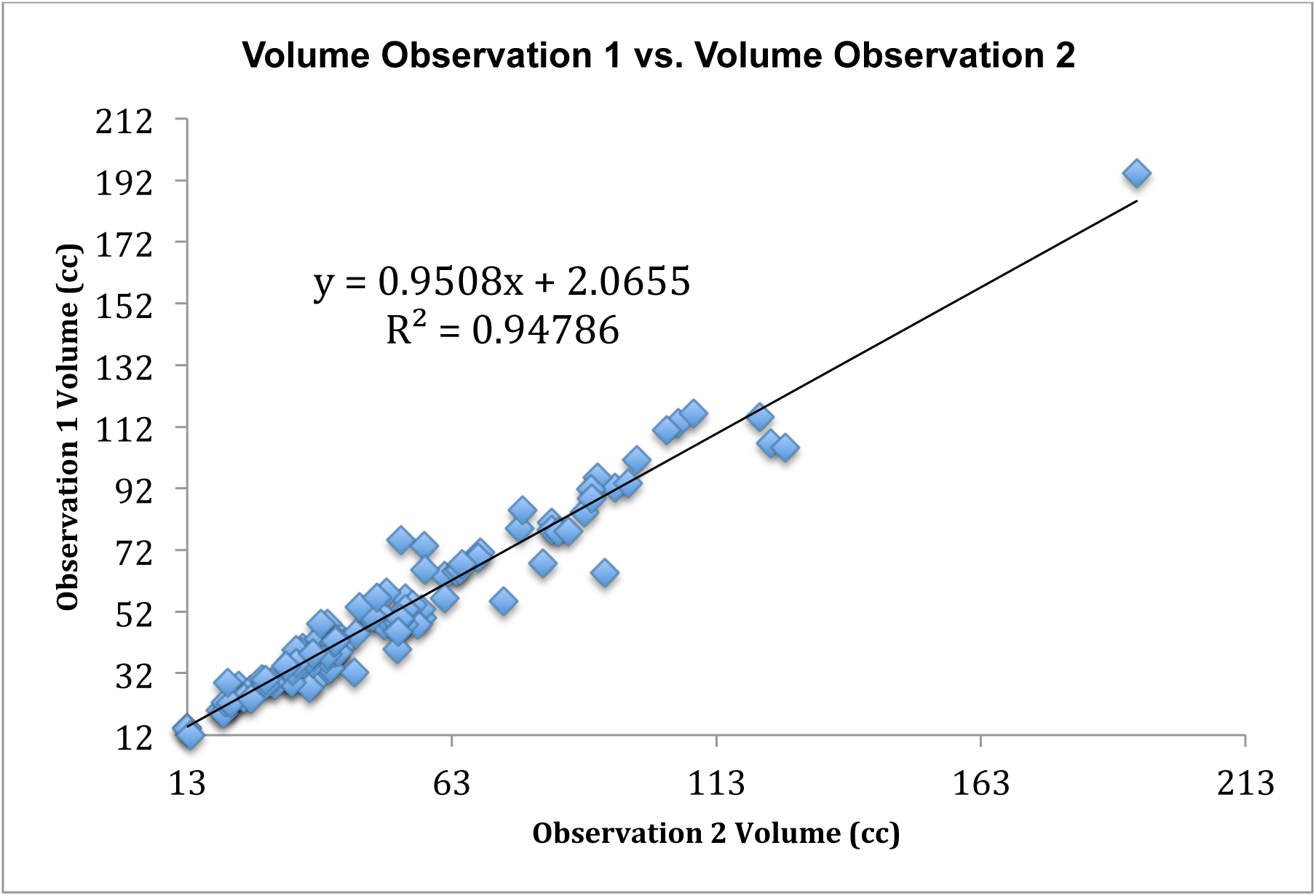
Intra-rater Concordance of Administrator

**Fig. 9A.**
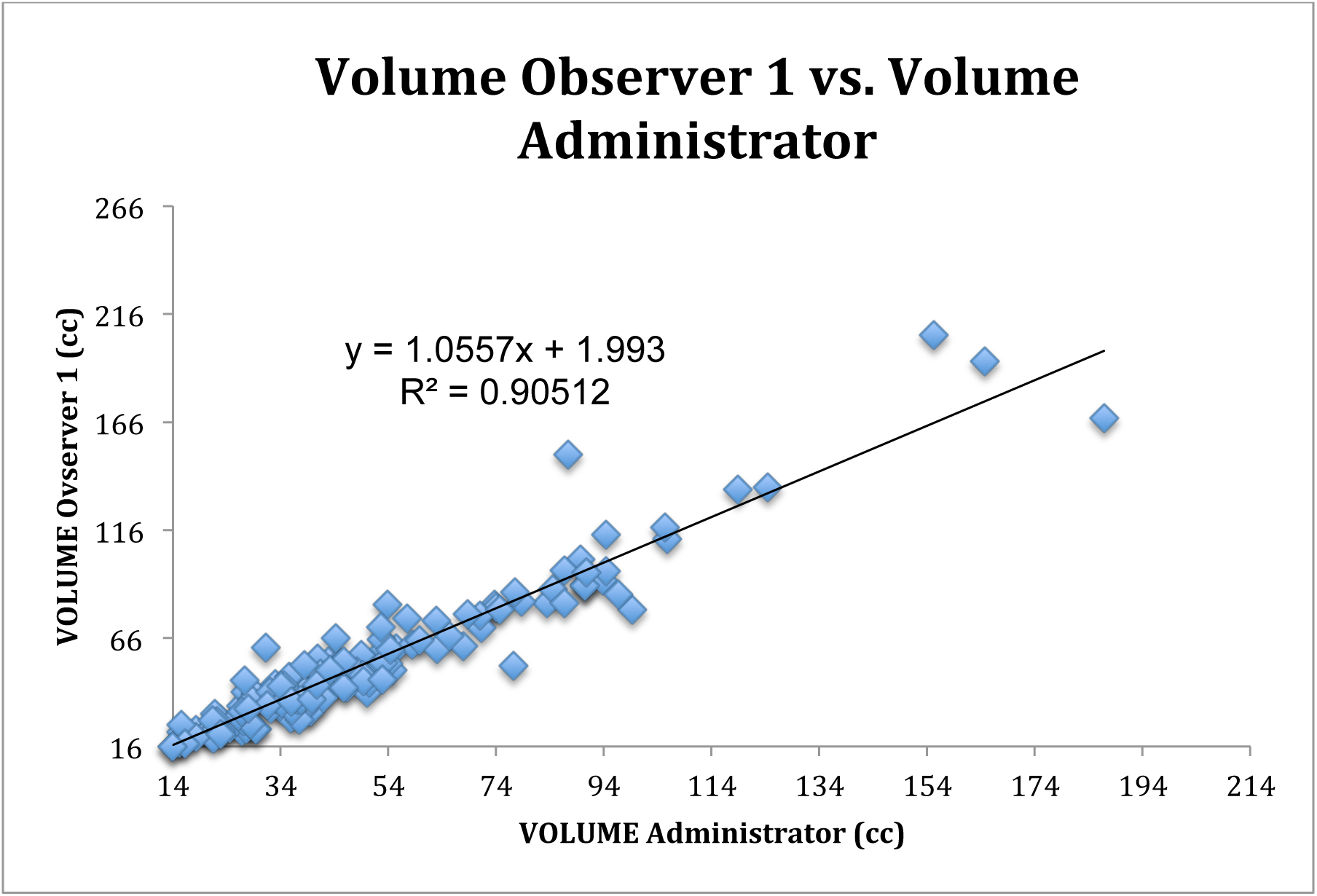
Accuracy Between Observer 1 and Administrator

**Fig. 9B.**
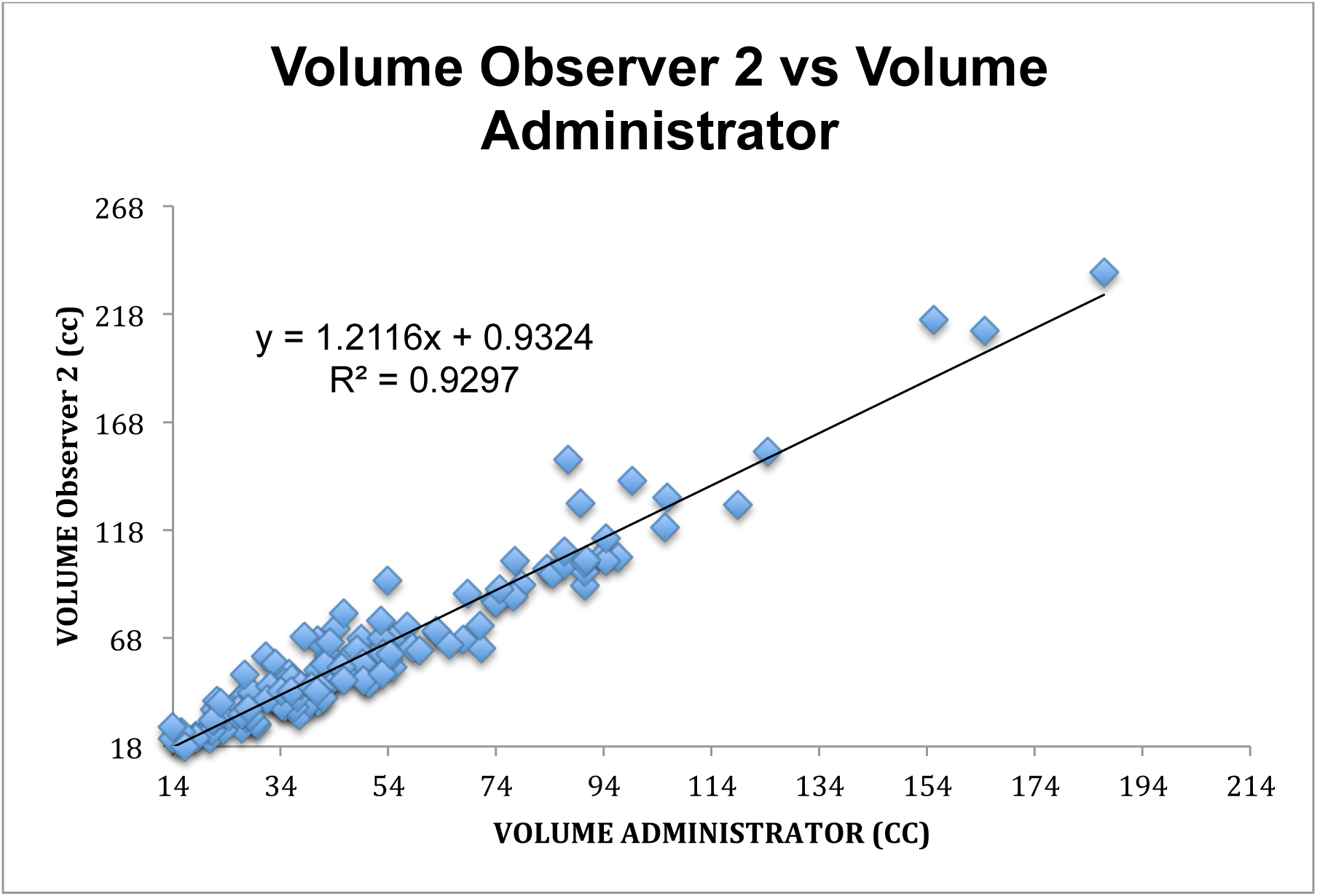
Interrater Concordance between Observer 2 and Administrator

**Fig. 10.**
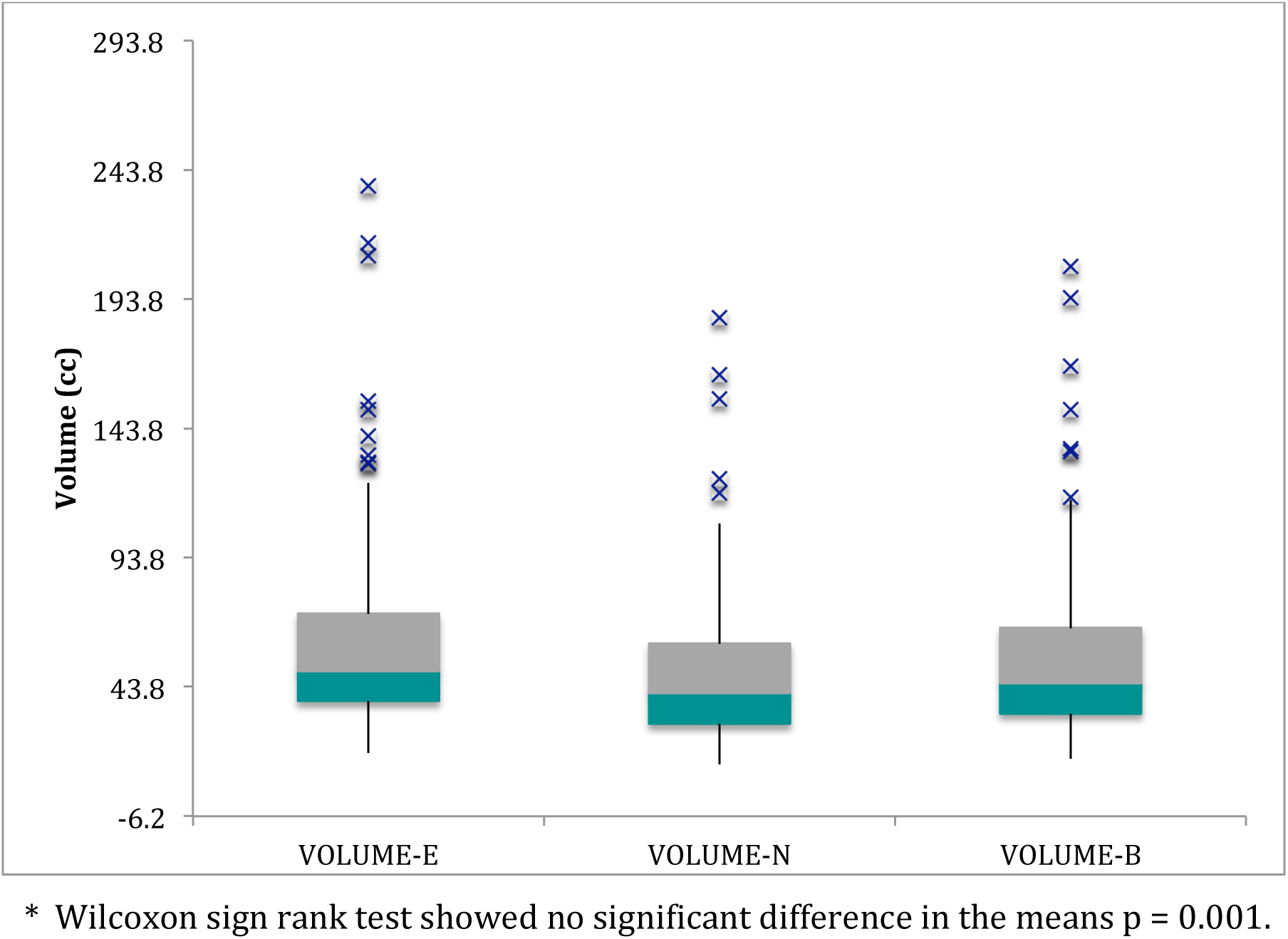
Box Plots of Prostate Volume Measurements of 2 Interpreting Radiologists (E and B) and Administrator (N)*

## Discussion

The purpose of this study was to determine the inter- and intra-rater reliability of an alternative method of measuring total prostatic volume based on well-defined anatomic landmarks in order to improve precision among observers. After review of the anatomy and correlation with MRI, the resultant measurements show excellent statistical agreement.

The conventional transverse measurement method attempts to find the outside EPC margins. Because of the adjacent position of pericapsular veins and in the mid- and lower levels the levator ani muscles, these structures are often included in the transverse measurement. MRI is superior to ultrasound in selecting the outer boundary of the EPC. We chose to select the inner boundary of the EPC for axial plane landmarks to improve interobserver consistency (precision). In the everyday clinical setting, measurement of the outer boundary of the EPC, when visible in the maximal transverse view, is acceptable. We also describe better criteria for identifying a more consistent selection of the anterior and posterior landmarks for measurement in the axial view. Some might take issue with our technique of not including the entire AFMS in some cases wherein the maximal transverse axial plane falls at an extreme proximal level. In these situations, the detrusor (primarily smooth muscle) component of the AFMS is focally thickest and not representative of the lower two-thirds of the prostate. In addition to the “salami” effect mentioned earlier, this may explain why EVF calculations in most studies indicate that volume in glands < 35 cc tend to be overestimated [12].

The single most common cause for variation of total prostatic volume between examiners is likely selection of precise lower and upper landmarks in the mid-sagittal plane while measuring length [5, 28]. Common errors include incorrect identification of the inferior boundary of the apex as the caudal landmark, inconsistent identification of the cranial landmark, making measurements in a parasagittal (rather than mid-sagittal) plane, and measuring from the anterior or posterior prostatic base to apex. Our understanding of the location of the inferior boundary of the PZ is critical. Experience has shown that relying only on the apparent lower margin of the PZ in the mid-sagittal plane can be misleading and that confirmation in the coronal view using a localizer function is required for accuracy. Collins, et al. [29] pointed out two methods of longitudinal methods of measurement, one using the bladder base as the cranial landmark and the other the superior margin of the prostatic tissue including any “median” lobe (Fig. 11). We refer to our proposed method of length measurement as “biproximate” as it conveniently embraces both.

**Fig. 11.**
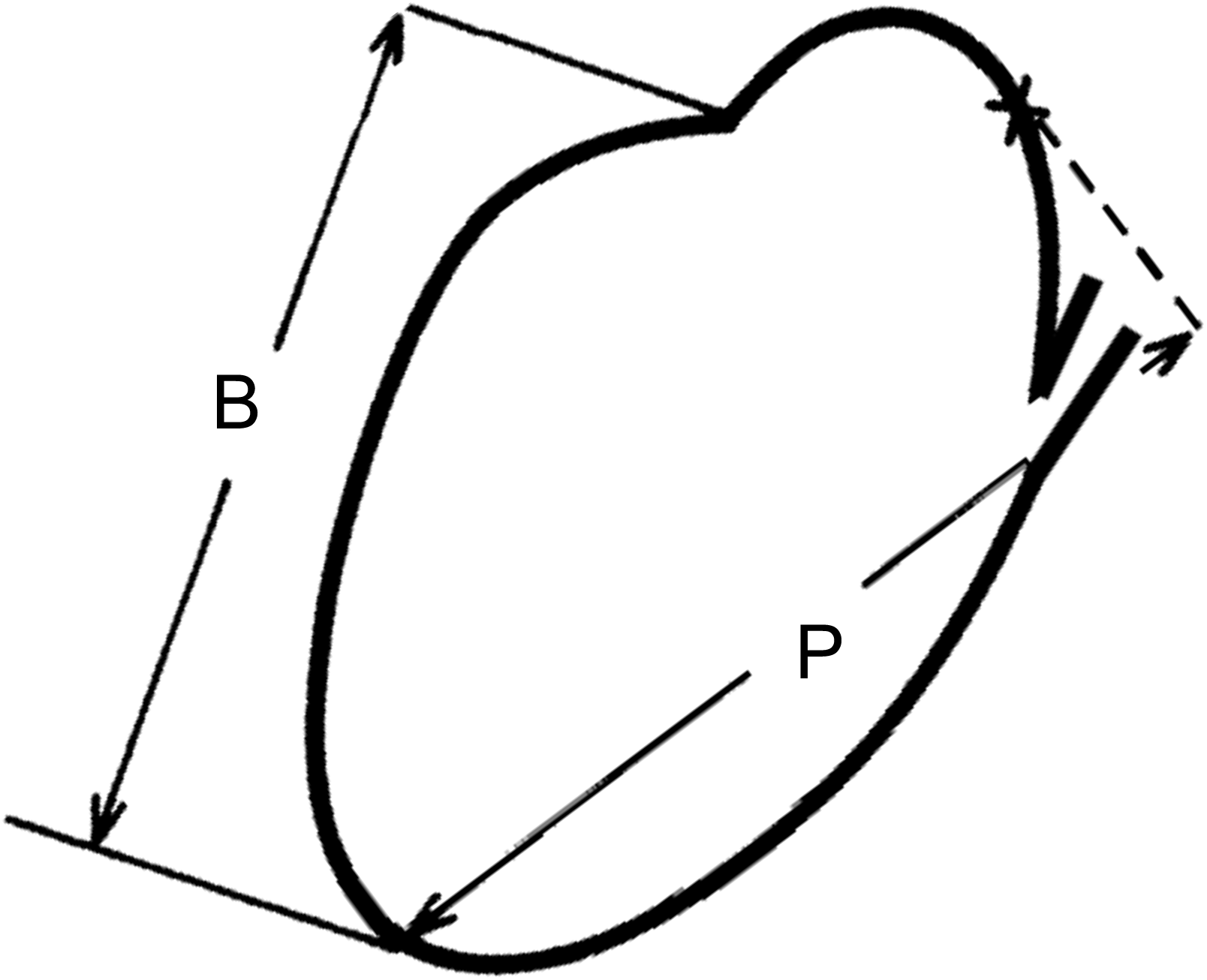
Two proposed ways to measure prostate length in the mid-sagittal plane. B=Distance from apex to bladder neck, PT= distance from apex to most proximal prostate tissue in sagittal plane (Modified from Collins GN, Ultrasound in Med. & Biol. 1995; 21:1102)

While measurement of the AP diameter of the RU may be useful for lobar classification of BPH [4], it is non-contributory in determining total prostatic volume. However, the proposed alternative method of volumetrics includes measurement of IPP that may be of use to urologists for patient management [30–39]. Some studies indicate that IPP may be an independent variable effecting surgical or medical outcome [40]. IPP of greater than 5 mm as measured on transabdominal ultrasound, has been associated with clinical signs and symptoms of bladder outlet obstruction [30, 31] and responsiveness to medical management [31, 32, 38].

Landmarks presented on MRI in our study are much more accurately visualized than on TRUS and could be expected to produce better intra- and interobserver concordance than those selected on TRUS. Another advantage of MRI, in contrast to transabdominal and TRUS, is that MRI does not require complete bladder distention for adequate visualization of landmarks.

The choice of an expert as proxy for “ground truth” measure of total prostatic volume rather pathological specimen may be considered a limitation or advantage of our study. Other clinical research studies have accepted the concept of expert consensus as a “gold standard” [42, 43]. One limitation of our study is that we used a single expert rather than a panel of experts due to the relative paucity of those with experience using the measurement techniques under examination. Expert opinion is also the current standard for evaluation of MRI prostate segmentation [44].

One could argue that since the objective should be to make *in vivo* measurements under normal physiological conditions, the volumes obtained were more realistic than under post-surgical or post-mortem conditions. *Ex vivo* measurements in the pathology department have disadvantages that reduce accuracy compared to those made in the living subject. The prostate is no longer infused with active circulation. This decreases specimen weight (volume) [15]. Processing of the specimen has been variable in previous studies. Comparison is usually made between the imaging measured volume and the “wet weight” of the fresh specimen on a scale or by water displacement in a graduated cylinder [45]. More accurate weights can be derived from electronic scales [14].

Since the specific gravity of prostate tissue and water are nearly identical (1.05 gm to 1.0 ml) [46], weight and volume have traditionally been used interchangeably in the literature, introducing mild inaccuracies when comparing image volumes to pathologic weights in larger glands. Some investigators use this number as a correction factor for comparing weight to volume [9, 47]. This assumption fails to take into account that it is likely that there is a spectrum of density values in larger prostates with different proportions of stromal, glandular, and malignant tissues [46]. There is no evidence that the presence of carcinoma changes the accuracy between TRUS and cadaver volumes [48]. The specimen may include the attached seminal vesicles, vas deferens and surrounding soft tissues in many studies [48]. In some, the seminal vesicles and vas deferens are physically removed before weighing [49, 14, 9], and in others the estimated mean volume of the seminal vesicles, as determined from the medical literature, is subtracted from the total weight of the prostate and accessory structures [15, 47]. Changes in prostate size due to drying or fixation complicate comparisons in cadaver studies [49, 50].

MRI volume imaging eliminates these variables, and it could be argued that MRI volumetrics in the living patient should replace post-mortem measurement as the “gold standard” [6]. Hass, et al. [51] has suggested that MRI-measured EVF might be considered the new standard for total volume estimate. In a small early small study using semi-automatic planigraphy, MRI showed improved results compared with TRUS [6, 9].

The application of artificial intelligence algorithms and the use of segmentation programs to establish prostate volume with MRI were reviewed by Turkbey, et al. [52]. These include innovative semi-automatic and automatic computer-based methods to “find” elusive prostate boundaries (edge detection). The software used is only available for research and not clinical use. These technologies share with geometric orthogonal models, such as the prolate ellipsoid formula (EVF) the fact that they are based on mathematical models and not direct measurement e.g. weight. The former requires significant boundary adjustments by an expert reader who still must ultimately understand the anatomy and landmarks we have described. At present, these techniques are thought by most to be too labor intensive for speedy clinical use [9, 47, 51]. Fully automatic unsupervised computerized assisted programs are under study, but performance compared with simpler methods is less well understood [44]. The great advantage of a fully automatic method would be precision since there is no user variation. Benxinque, et al [53] recently compared prolate ellipsoid and other prostate volume measures to a commercially available fully automated segmentation program. Although fully automatic measurement was not reliable compared to manually adjusted MRI segmentation, they demonstrated strong correlation of manually adjusted MRI 3D segmentation by an experienced radiologist relative to a medical student (ICC 0.91) and relative to MRI EVF (ICC 0.90). They also found that in the hands of a radiologist, the manually adjustment time was acceptably short.

It should be remembered that prostate imaging is not necessary in the initial work-up of lower urinary tract symptoms, and only is considered when surgical or minimally invasive procedures are under discussion [54]. TRUS, because of its cost effectiveness and availability will likely continue to be the basic clinical volumetric tool. But MRI will be useful for research studies of treatments requiring follow-up monitoring [6] and in the clinical assessment of larger prostates and could replace ultrasound if limited protocols can be found and costs reduced.

It may be time to accept MRI as the new “gold standard” for assessing prostate volume, especially in research protocols [6, 12, 51].

## Conclusions

We present a succinct review of the scientific foundations for identifying the anatomic boundaries necessary for reproducible multiplane volume measurement of the prostate using MRI. Detailed landmarks for these measures are sufficiently shown to allow for reliable transfer of method to the general community of radiologists and urologists. An alternative method of measurement (biproximate) is introduced to enhance measurement consistency and to answer the needs of operating urologic surgeons interested in the effects of IPP. The measurements tested showed excellent inter- and intraobserver reliability and is suitable especially for multi-observer research.

*The authors declare no competing interests. All original data is available by email from the lead author (NFW) upon reasonable request*.

## Author Contribution Statement

NFW: Conceptual idea, prostate anatomy, experimental design, data collection and assessment, statistical calculations, writing of primary manuscript

EN: Data collection and Editorial review and critique

BS: Data collection and Editorial review and critique

## Compliance and Ethics Statement

The Internal Review Board for research and Ethics Committee are combined at the University of Minnesota into a single on-line submission called Ethosirb for which there is a single identifying protocol number (00003941). Informed consent was waved for this retrospective study carried out in accordance with the relevant guidelines and regulations of our institutional Internal Review Board and Ethics Committee, meeting all international standards for human research.

